# Computational Discovery of CRISPR-Cas13b Guide RNAs for Broad-Spectrum Dengue Virus Targeting

**DOI:** 10.64898/2025.12.31.697158

**Authors:** Syed Muhammad Ali Naqvi, Farhan Khan, Muslim Hussain

**Affiliations:** Computer Science, Habib University, Karachi, Pakistan; Electrical and Computer Engineering, Habib University, Karachi, Pakistan

**Keywords:** CRISPR-Cas13b, Dengue Virus, Machine Learning, Guide RNA Design, Antiviral Therapy, Computational Biology

## Abstract

Dengue (DENV), an RNA virus, remains a significant global health threat, particularly in developing regions, with no widely effective antiviral therapy available. The CRISPRCas13b system, specifically the *PspCas13b* subtype, has emerged as a promising programmable antiviral tool capable of targeting viral RNA with high specificity. However, the efficacy of Cas13b-based interventions relies heavily on the design of potent and conserved CRISPR RNA (crRNA) spacer sequences, a task complicated by high viral genetic diversity. Unlike CRISPR-Cas9, which targets double-stranded DNA in eukaryotic genomes, Cas13b directly targets single-stranded RNA, making it ideally suited for RNA virus therapeutics; however, existing computational tools predominantly focus on Cas9 DNA targeting or Cas13d for mammalian transcript knockdown, leaving a significant gap for Cas13b-specific viral antiviral design.

In this paper, we propose a computational pipeline and machine learning framework for the rational design of high-efficacy Cas13b guide RNAs targeting all four Dengue serotypes. Our approach integrates large-scale genomic data extraction, conservation analysis, and a novel *in silico* optimization module for guide RNA (gRNA) sequences, based on recently reported Cas13b design rules (e.g., 5’ GG motif preference, Cytosine penalties). To predict targeting efficiency, we benchmark classical machine learning models (Random Forest, XGBoost) against foundation model-based predictors (Nucleotide Transformer, RNA-FM) using a dataset of experimentally validated spacers.

Our results demonstrate that classical feature-engineered models significantly outperform deep learning approaches when trained on experimentally validated gRNA datasets in low-data regimes. We identify highly conserved, optimized crRNA candidates, including several pan-serotype guides with predicted high potency. This work establishes a baseline for Cas13b efficiency prediction and provides a robust computational resource for accelerating the development of CRISPR-based antivirals against Dengue and other RNA viruses.

Code: https://github.com/muhammadali74/CAS13b_pipeline

## I. Introduction

Dengue Virus (DENV) remains one of the most critical mosquito-borne viral pathogens worldwide, posing a severe public health burden, particularly in developing nations such as Pakistan, India, and Brazil [1], [2]. With four distinct serotypes (DENV-1 through DENV-4), the virus causes millions of infections annually, ranging from mild fever to fatal hemorrhagic fever. Despite the prevalence of the disease, therapeutic options remain limited. Developing effective antivirals or vaccines is complicated by the high genetic diversity of the virus and the phenomenon of antibody-dependent enhancement (ADE), where sub-optimal antibodies can worsen symptoms upon secondary infection [3]. Consequently, there is an urgent need for precise, molecular-level interventions that can target conserved viral regions across all serotypes.

The CRISPR-Cas13 system has emerged as a revolutionary tool for RNA targeting. Unlike the DNA-targeting Cas9, Cas13 enzymes (such as Cas13a, Cas13b, and Cas13d) target single-stranded RNA, making them ideal candidates for combating RNA viruses like Dengue. Among these, the Cas13b subtype (specifically *PspCas13b*) has demonstrated high specificity and minimal off-target effects, rendering it a promising candidate for therapeutic and diagnostic applications [4]. However, the efficacy of Cas13b is highly dependent on the design of the CRISPR RNA (crRNA) spacer sequence. Selecting a 30-nucleotide (nt) spacer that is both conserved across viral mutations and highly efficient in triggering Cas13b activity is a multi-objective optimization problem that is computationally non-trivial.

Traditional selection of guide RNAs often relies on simple sequence alignment or tedious manual screening in the wet lab, which is cost-prohibitive and time-consuming. While Artificial Intelligence (AI) and Machine Learning (ML) have been successfully applied to Cas9 and Cas13d guide design [5]–[7], there is a distinct lack of computational frameworks specifically tailored for Cas13b, particularly models trained to navigate the constraints of viral conservation.

In this paper, we present an end-to-end computational pipeline and a baseline Machine Learning framework for the rational design of Cas13b crRNAs against Dengue Virus. Our approach integrates genomic data mining, conservation analysis, and predictive modeling to identify potent viral targets. We leverage recent findings regarding Cas13b’s specific nucleotide preferences to optimize these spacers in silico. Furthermore, we address the challenge of limited experimental data by benchmarking classical ML models against large language model (LLM) embeddings (such as Nucleotide Transformer and RNA-FM), establishing a robust baseline for Cas13b efficiency prediction.

The main contributions of this work are as follows:

- A comprehensive data processing pipeline that identifies highly conserved 30-nt spacers across all four Dengue serotypes from public genomic databases.
- A novel optimization module that enhances crRNA efficacy by introducing strategic mismatches based on specific Cas13b design rules.
- A comparative analysis of ML models for predicting Cas13b target efficiency, demonstrating the effectiveness of feature engineering over deep learning embeddings on small-sample datasets.
- A catalog of top-ranked, optimized crRNA candidates suitable for experimental validation.

## II. Literature Review

### A. CRISPR-Cas13 as an Antiviral Strategy

The discovery of CRISPR-Cas systems adapted for RNA targeting has opened new avenues for antiviral therapy. While Cas9 targets DNA, the Class 2 Type VI CRISPR systems, characterized by the effector protein Cas13, possess RNase activity capable of cleaving single-stranded RNA [8]. Early applications utilized Cas13a (formerly C2c2) for pathogen detection, such as the SHERLOCK platform. More recently, focus has shifted toward Cas13b and Cas13d for therapeutic silencing due to their lack of protospacer flanking site (PFS) requirements in eukaryotic cells [9]. In the context of RNA viruses, Cas13 has been shown to effectively inhibit Influenza A, SARS-CoV-2, and Dengue in cell cultures, positioning it as a programmable antiviral agent [10].

### B. Design Rules for PspCas13b

For a CRISPR system to be effective, the guide RNA must hybridize efficiently with the target. While Cas13 systems generally exhibit tolerance to mismatches, recent studies have elucidated specific design constraints for *PspCas13b*. Hu et al. [11] conducted a systematic screening of Cas13b crRNAs, revealing that secondary structure has a moderate effect on efficacy, while specific nucleotide motifs drive potency. Specifically, a “GG” motif at the 5’ end of the spacer (positions 1-2) typically enhances activity, whereas Cytosine (C) residues at positions 1-4 and the central region (11-17) act as negative predictors. These findings imply that simply selecting guide RNAs based on conservation of the target viral sequence is insufficient; a pipeline must optimize sequences to satisfy these biochemical preferences, even if it requires introducing artificial mismatches, which Cas13b is known to tolerate [11].

### C. Machine Learning in CRISPR Guide Design

The application of Machine Learning to predicting CRISPR guide efficiency is well-established for Cas9. Tools such as DeepCRISPR [5] and CRISPR-scan [12] utilize deep neural networks and extensive feature engineering to predict on-target efficiency and off-target risks. However, these models are not transferable to Cas13 due to fundamental differences in the cleavage mechanism and target substrate (RNA vs. DNA).

For Cas13 systems, recent tools like CASowary [13], and prediction models for Cas13d [14] [6] have begun to emerge. These typically rely on Convolutional Neural Networks (CNNs) trained on high-throughput screening data. However, for Cas13b specifically, public datasets remain scarce. The challenge is further compounded when applying large genomic foundation models. While Transformer-based models like Nucleotide Transformer [15] and RNA-FM [16] excel at capturing long-range genomic dependencies, their application to regression tasks on small, specialized datasets (such as Cas13b efficiency data) often results in overfitting or poor generalization compared to classical ensemble methods using domain-specific engineered features. This paper addresses this gap by developing a specialized pipeline that combines bio-informatic filtering with an ML model optimized for the specific constraints of Cas13b.

## III. Methodology

Our proposed pipeline integrates bioinformatics filtering, conservation analysis, sequence optimization, and machine learning-based efficacy prediction to identify high-potency Cas13b crRNA candidates for Dengue Virus. The workflow consists of five main stages, as follows.

### A. Data Acquisition and Preprocessing

Dengue Virus genome sequences were retrieved from the Bacterial and Viral Bioinformatics Resource Center (BV-BRC) database. The raw dataset comprised genomic sequences spanning all four Dengue serotypes (DENV-1 through DENV-4) as well as unclassified strains. To ensure data quality, sequences were filtered based on two criteria:

- **Ambiguity threshold:** Sequences containing more than 5% ambiguous nucleotides (denoted as ‘N’) were excluded to prevent spurious spacer extraction.
- **Minimum length:** Only genomes exceeding 1,000 base pairs were retained, as shorter fragments are unlikely to represent complete viral coding regions.

Each genome was annotated with its corresponding serotype by parsing the genome name metadata. The final filtered dataset was converted into FASTA format for downstream processing. Post-filtering statistics are presented in Table I.

**TABLE I.**
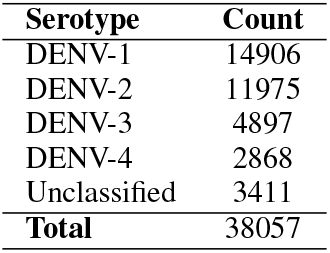
Genome Statistics After Quality Filtering.

### B. Spacer Extraction and Conservation Analysis

To identify candidate crRNA spacers, we employed a sliding window approach to extract all possible 30-nucleotide (nt) subsequences from each viral genome. A memory-efficient SQLite-based storage system was implemented to handle the large combinatorial space of candidate spacers across thousands of genomes.

For each unique 30-nt spacer:

- We tracked the set of genome identifiers in which it appeared, avoiding redundant counting within the same genome.
- Conservation percentage was computed as

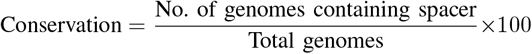
- Subtype-specific occurrence counts (DENV-1, DENV-2, DENV-3, DENV-4, Unclassified) were calculated by mapping genome IDs to their serotype annotations.

This process yielded millions of unique spacers. To focus on the most promising candidates, we selected the top-300 most frequently occurring spacers per serotype, ensuring broad coverage across all viral variants.

### C. Sequence Filtering and Quality Control

Each candidate spacer was annotated with biochemical properties known to influence Cas13b activity:

- **GC Content:** Computed as the percentage of Guanine and Cytosine nucleotides. Extreme GC values can affect transcription and secondary structure stability.
- **Homopolymer Runs:** Binary flag indicating the presence of four or more consecutive identical nucleotides, which can cause transcriptional termination or synthesis artifacts.

While these features were computed for all spacers, strict filtering was not applied at this stage, as subsequent optimization and machine learning prediction would further refine candidate selection.

### D. Cas13b-Specific Sequence Optimization

Recent functional studies [11] have revealed that PspCas13b activity is strongly influenced by position-specific nucleotide preferences. We adapted the scoring metric the authors used for our optimization algorithm given as follows:

- A Guanine (G) at positions 1 or 2 (5’ end) confers a score bonus of +20 each.
- Cytosine (C) at positions 1–4 incurs a penalty of −20 each.
- Cytosine at positions 11, 12, 15, 16, and 17 incurs a penalty of −5 each.

Based on this scoring scheme, we developed an optimization algorithm that introduces up to three strategic nucleotide substitutions to maximize the Cas13b potency score while maintaining biological relevance. The algorithm operates as follows:

1) Compute the baseline score for the original spacer.

2) Identify all positions where beneficial mutations exist (e.g., replacing C with G at penalized positions).

3) Generate all combinations of 1, 2, or 3 mutations.

4) Score each variant and retain those achieving the maximum score.

5) If multiple variants share the top score, all are preserved for downstream analysis.

This strategy leverages the known mismatch tolerance of Cas13b [11], which can accommodate up to 3–4 nonconsecutive mismatches without significant loss of activity.

### E. Machine Learning Model for Efficiency Prediction

1. *Training Dataset:* A curated dataset of 245 Cas13b spacer sequences with experimentally validated mean targeting efficiencies was used for model training. The data were extracted from the supplementary data of [11], and the specific source table, along with the spacers used in this study, is provided in Table IV. This dataset represents high-throughput screening results from previous studies.
2. *Feature Engineering:* Given the limited sample size (n=245), we employed extensive feature engineering to encode domain-specific knowledge:
  - **Position-Specific One-Hot Encoding:** Each of the 30 nucleotide positions was encoded as a 4-dimensional binary vector (A, T, G, C), yielding 120 binary features.
  - **Motif Features:** Explicit Boolean features for critical motifs (e.g., ‘GG’ at positions 1–2, C-count at positions 1–4).
  - **Sequence Composition:** GC content, nucleotide fractions, and di-nucleotide frequencies. A complete list of the 157 engineered features is provided in the Table V in supplementary materials.
3. *Model Selection and Training:* We compared multiple regression models suitable for small, high-dimensional datasets:
  - Random Forest Regressor
  - Gradient Boosting Regressor
  - XGBoost Regressor
  - Support Vector Regressor (SVR)
  - Ridge Regression Hyperparameter tuning was performed using 5-fold cross-validation with grid search, optimizing for *R*^2^ and minimizing overfitting. The final model was trained on all 245 samples and saved for inference.
4. *Deep Learning Experiments:* To explore whether large pre-trained genomic foundation models could improve upon classical ML approaches, we conducted two additional experiments using state-of-the-art sequence embeddings. The motivation for this approach stems from the success of transformer-based language models like BERT and GPT) in NLP tasks and their recent adaptation to genomic sequences [15], [16]. Such models, trained on billions of nucleotides, may capture complex sequence patterns and biophysical properties that are difficult to encode manually. We hypothesized that these learned representations could provide richer feature spaces than hand-crafted features, potentially improving predictive power for Cas13b efficiency.

We experimented with two foundation models:

1. **Nucleotide Transformer (NT):** A BERT-style transformer pre-trained on genomic DNA sequences was used to extract 512-dimensional embeddings for each spacer. A lightweight regression head (2-layer MLP) was trained on these embeddings.
2. **RNA-FM:** A foundation model pre-trained on RNA sequences was similarly used for embedding extraction, followed by regression head training.

Both approaches were trained with heavy regularization (dropout=0.3, weight decay=0.01) and early stopping to mitigate overfitting.

### F. Candidate Ranking and Selection

The trained machine learning model was applied to all optimized spacer sequences. Candidates were ranked by:

1. Predicted targeting efficiency
2. Conservation across serotypes
3. Cas13b position score

For each serotype, we identified:

- The top-3 most conserved spacers (broad-spectrum candidates)
- All spacers with predicted efficiency≥ 0.8 (high-potency candidates)

## IV. Experiments and Results

### A. Spacer Extraction and Conservation

The sliding window extraction across all filtered genomes yielded 3.78 million unique 30-nt spacers. After subtype-specific conservation ranking and selection of the top-300 per serotype, we retained 1740 candidates for optimization.

### B. Sequence Optimization Results

The Cas13b-specific optimization algorithm significantly improved spacer quality. Of the 1,740 input spacers, 1,714 sequences (98.5%) received at least one beneficial mutation, while 26 sequences (1.5%) were already optimal according to the scoring scheme.

The optimization yielded substantial improvements in predicted potency:

- **Average Score Improvement:** +47.17 points (SD = 20.37)
- **Maximum Score Improvement:** +100 points

The majority of spacers required multiple mutations to achieve maximum potency: 9.9% required one mutation, 23.7% required two mutations, and 64.9% required all three allowed mutations. Score improvement correlated monotonically with mutation count, yielding average improvements of 17.8, 40.2, and 55.6 points for one, two, and three mutations, respectively.

Position-specific analysis revealed that 73.3% of all mutations occurred at positions 1–2 (introducing the favorable ‘GG’ motif), while the remaining mutations targeted penalized Cytosine residues in positions 3–4 and the central region (11– 12, 15–17). Notably, conservation percentage did not predict optimization potential: highly conserved spacers (*≥*80%) benefited equally from optimization as less conserved candidates, indicating that sequence conservation and biochemical potency represent orthogonal selection criteria. Figure 1 illustrates these optimization dynamics across all dimensions of the analysis.

**Fig. 1.**
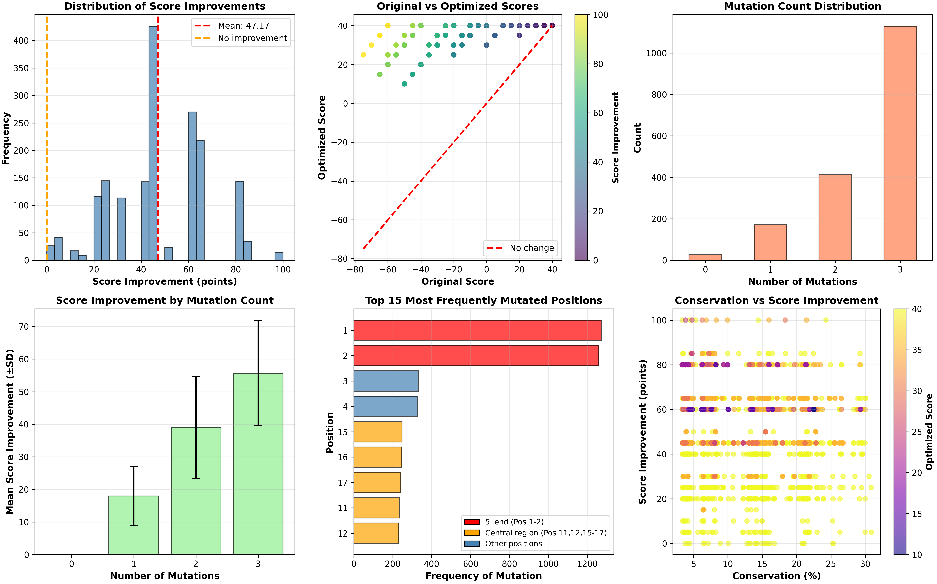
Comprehensive analysis of Cas13b-specific sequence optimization. (a) Distribution of score improvements showing mean of +47.17 points. (b) Scatter plot of original vs optimized scores colored by improvement magnitude. (c) Distribution of mutation counts per spacer. (d) Mean score improvement stratified by mutation count. (e) Top 15 most frequently mutated positions, with red indicating 5’ end (positions 1–2) and orange indicating central penalty region (positions 11–12, 15–17). (f) Conservation vs score improvement, demonstrating orthogonal relationship between sequence conservation and optimization potential.

### C. Machine Learning Model Performance

1. *Classical ML Models:* Table II summarizes the 5-fold cross-validation performance of all tested models. **Best Model:** XGBoost achieved the highest cross-validated *R*^2^ of 0.235, a lowest MAE of 0.161 and a high Spearman correlation of 0.529, indicating moderate predictive power suitable for ranking candidate spacers.
2. *Deep Learning Model Performance:* Despite leveraging large pre-trained models, the deep learning approaches underperformed relative to classical ML. This is most likely due to the insufficient data needed for training a deep-learning models

- **Nucleotide Transformer (NT-DL):**
  – Mean *R*^2^: 0.0181
  – Mean Spearman: 0.2607

- **RNA-FM:**
  – Mean *R*^2^: 0.1131
  – Mean Spearman: 0.3429

**TABLE II.** Machine Learning Model Comparison (5-Fold CV)

| Model | Mean $R^2$ | Mean MAE | Mean Spearman |
| --- | --- | --- | --- |
| XGBoost | 0.235 | 0.161 | 0.529 |
| Gradient Boosting | 0.199 | 0.220 | 0.497 |
| Random Forest | 0.192 | 0.221 | 0.480 |
| SVR | 0.227 | 0.217 | 0.533 |
| Ridge Regression | 0.139 | 0.231 | 0.494 |

RNA-FM showed modest improvement over NT, likely due to its RNA-specific pre-training, but both failed to match the engineered-feature baseline. This outcome is consistent with the literature: foundation models require substantial task-specific data to outperform domain-engineered features in low-data regimes [15], [17], [18].

### D. Feature Importance Analysis

SHAP (SHapley Additive exPlanations) analysis was performed on the best-performing model to interpret feature contributions. The analysis revealed that Cas13b efficiency is driven by a hierarchy of sequence composition features rather than single-position identities alone. The most influential features (see Figure 2) were:

**Fig. 2.**
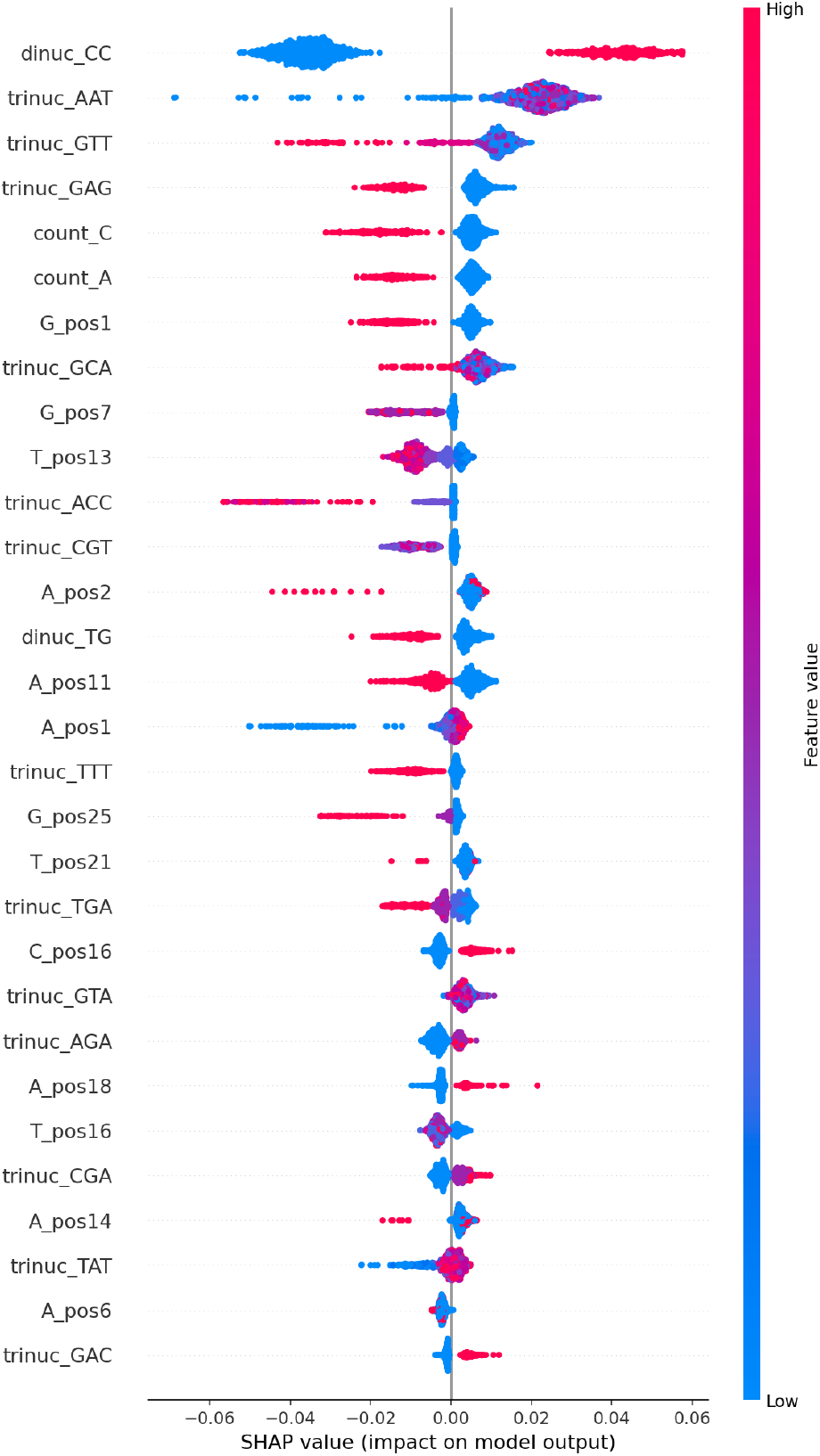
SHAP summary plot ranking the top 30 features by mean absolute SHAP value. Red dots indicate high feature values, while blue dots indicate low values. The x-axis shows the impact on predicted Cas13b efficiency (positive = higher efficiency). The plot reveals that dinucleotide CC frequency (`dinuc_CC`) is the single most dominant predictor, strongly negatively correlated with efficiency, aligning with known Cas13b preference against Cytosine-rich motifs.

**1) CC dinucleotide frequency** `(dinuc_CC)`: This was the single most dominant predictor. High values (red dots) had a massive negative impact on predicted efficiency (SHAP value *≤*-0.04), confirming the strong deleterious effect of Cytosine enrichment.

**2) Trinucleotide context:** Specific trimers such as `AAT, GTT,` and `GAG` were highly predictive, suggesting that local sequence context significantly modulates activity.

**3) Overall Cytosine count** `(count_C)`: Consistently showed a negative correlation with efficacy, reinforcing the rule that C-rich spacers perform poorly.

**4) G at Position 1** `(G_pos1)`: Confirmed as a top positive predictor, validating the 5’ G preference identified in prior literature.

**5) Specific positional nucleotides:** Features such as `G_pos7, T_pos13,` and `A_pos2` appeared as key modulators, indicating that positions beyond the canonical 5’ end also contribute to spacer potency.

These findings align with and extend the experimental results from Hu et al. [11]. While previous studies emphasized specific penalty positions, our model learned that global Cytosine content (captured by `dinuc_CC` and `count_C`) is the primary determinant of low efficiency, providing a broader heuristic for guide design.

### E. Final Candidate crRNAs

Post-optimization, we filtered the 1,740 unique spacers to identify the top-performing candidates specific to each Dengue serotype. This selection process yielded between 269 and 396 high-confidence candidates per serotype (approximately 15– 22% of the total pool), prioritizing those with the highest frequency of occurrence in their respective genomic populations.

Figure 3 illustrates the relationship between conservation coverage and predicted targeting efficiency. We observed distinct landscape differences across serotypes:

- **DENV-3 and DENV-4 (High Convergence, Limited Representation):** Although these serotypes exhibited the most promising dual-purpose candidates with simultaneous high conservation and efficiency, they were underrepresented in the dataset. DENV-3 and DENV-4 accounted for 22.8% and 21.5% of the optimized candidates, respectively, compared to the 20% representation of larger serotypes. For DENV-4, the most conserved spacer `(GGAACA…)` covers 80.51% of all available genomes while maintaining a high predicted efficiency of 0.836. Similarly, the top DENV-3 candidate covers 69.88% of genomes with a robust efficiency of 0.859. These candidates demonstrate ideal therapeutic potential but would benefit from validation against a larger, more diverse strain collection.
- **DENV-1 (Balanced Representation):** With 349 candidates (20.1% of the pool), DENV-1 showed a dense cluster of high-efficiency spacers across moderate conservation ranges. The most conserved candidate covers 68.42% of genomes with a predicted efficiency of 0.804, offering a strong balance for clinical application. The highest-efficiency candidate achieves 0.881, demonstrating that DENV-1 spacers can be optimized for either criterion without severe trade-offs.
- **DENV-2 (Trade-off with Compensatory Options):** DENV-2 (352 candidates, 20.2% of pool) presented a notable divergence between conservation and potency. The most conserved spacer (68.75% coverage) had a markedly lower predicted efficiency of 0.613. However, the highest-efficiency candidate `(GGAGAC`…, efficiency 0.878) covers only 42.36% of genomes. Importantly, subsequent high-ranking candidates offer a viable middle ground: the fourth-ranked candidate achieves a mean targeting efficiency of 0.832 while maintaining reasonable coverage, presenting a practical compromise for serotype-specific applications. For maximal pan-serotype coverage, we recommend either the efficiency-optimized guide or a multi-guide strategy incorporating both high-efficiency and high-conservation targets.

**Fig. 3.**
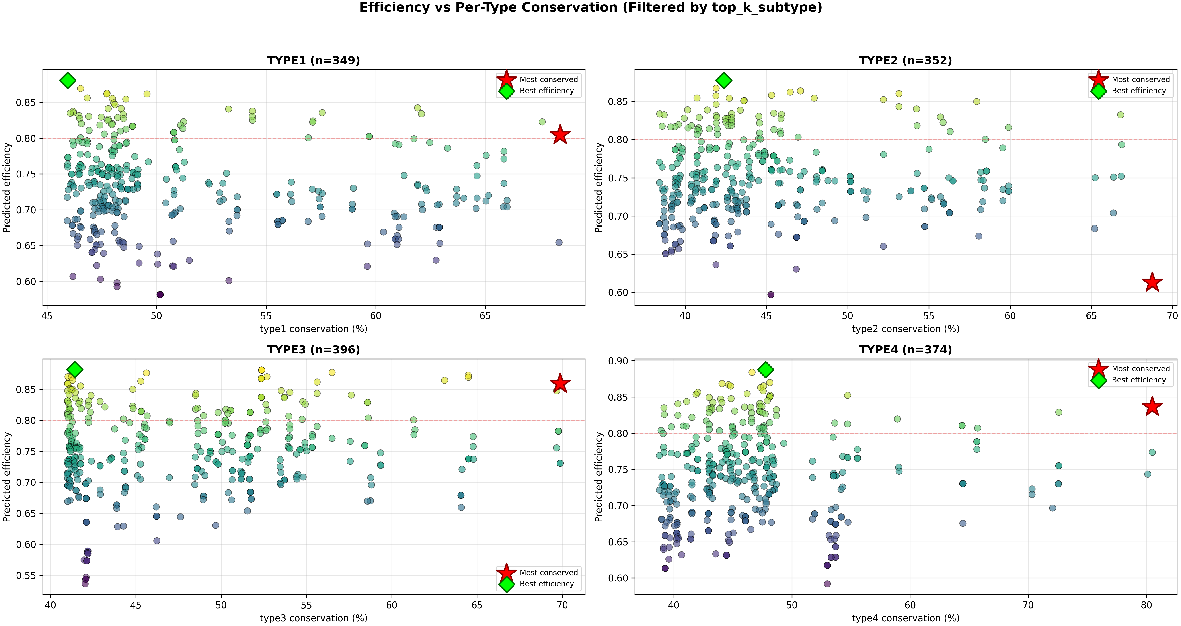
Predicted efficiency versus serotype-specific conservation for the top candidate crRNAs. Each point represents a unique spacer optimized for DENV-1 (*n* = 349), DENV-2 (*n* = 352), DENV-3 (*n* = 396), and DENV-4 (*n* = 374). The **Green Diamond** marks the candidate with the highest predicted efficiency, while the **Red Star** marks the candidate with the highest conservation coverage for that serotype. The red dashed line indicates the high-efficiency threshold (*≥* 0.8).

Table III lists the specific sequences for the optimal candidates identified in Figure 3. More candidate rankings are provided in Table VI.

**TABLE III.**
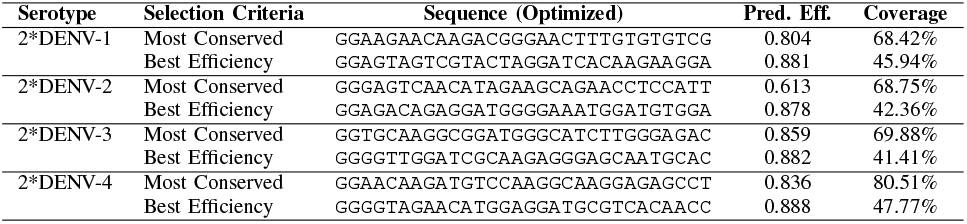
Best Candidate crRNAs per Serotype.

## V. Discussion and Analysis

### A. Biological Significance of Identified Candidates

The computational pipeline successfully identified serotype-specific crRNA spacers with distinct biological properties tailored to Cas13b targeting:

- **Highly conserved targets:** Most candidates maintain *≥* 45% representation within their respective serotype genomes, with optimal candidates reaching 68% coverage. This high conservation reduces the likelihood of viral escape through mutation and suggests that targeted regions encode functionally constrained viral elements (likely essential for replication or virulence).
- **Optimized for Cas13b biochemistry:** The extensive optimization phase (yielding 98.5% improvement rate with mean +47.17 points) demonstrates that even moderately suboptimal natural sequences can be substantially enhanced through strategic nucleotide substitutions. The algorithm-generated guides exploit known Cas13b preferences (5’ GG motifs, central C-elimination) while maintaining high specificity through the model’s learned features, which prioritize position-specific context and dinucleotide composition over simple global metrics.
- **Predicted to be highly effective:** Machine learning predictions identified top candidates. These predictions were informed by 158 engineered features validated against a curated dataset of 245 experimentally characterized Cas13b spacers, providing robust estimates grounded in observed biochemical principles.

### B. Performance of Classical ML vs. Deep Learning

Our results demonstrate that in data-scarce regimes (n=245), **feature-engineered classical ML models significantly outperform deep learning approaches**. This finding has important implications:

- **Sample efficiency:** Random Forest and XGBoost effectively leverage domain knowledge encoded in features, whereas foundation models (NT, RNA-FM) rely on emergent patterns from large-scale pre-training that may not align with task-specific objectives.
- **Interpretability:** SHAP analysis of tree-based models directly reveals which biochemical properties drive predictions, facilitating biological insight and model refinement.
- **Future potential:** As larger Cas13b efficiency datasets become available (e.g., from high-throughput screens), fine-tuning foundation models may become competitive. Our negative results establish a baseline for comparison.

### C. Comparison with Existing Guide Design Tools

To our knowledge, this is the **first computational framework specifically tailored for Cas13b crRNA design against viral genomes**. Existing tools like CASowary [13] target Cas13a/d for mammalian transcript knockdown, while platforms such as CHOPCHOP [19] focus on Cas9. Our pipeline addresses the unique constraints of antiviral applications:

- Conservation analysis across viral serotypes (not applicable to fixed genomes)
- Mismatch tolerance and optimization (viral diversity)
- Integration of virus-specific and enzyme-specific design rules

### D. Limitations and Future Work

- **Experimental validation:** The predicted candidates require wet-lab testing to confirm efficacy in Dengue-infected cells.
- **Off-target analysis:** While our spacers are specific to Dengue genomes, computational screening against the human transcriptome is recommended before therapeutic application.
- **Expanded training data:** Acquiring more Cas13b efficiency data (especially for viral targets) would enable more robust ML models and potentially unlock the advantages of foundation models.
- **Delivery mechanisms:** Translation to therapy requires development of efficient Cas13b delivery systems, which is beyond the scope of this work.

## VI. Conclusion

We have presented a comprehensive computational pipeline for the rational design of Cas13b crRNAs targeting Dengue Virus, addressing a critical gap in antiviral therapeutic development. By integrating genomic data extraction, sequence optimization, and machine learning-based efficacy prediction, we identified a catalog of high-confidence candidate guide RNAs suitable for immediate experimental validation. Our work establishes the first baseline for Cas13b efficiency prediction using classical machine learning, demonstrating superior performance over large language model embeddings in small-data settings. The identified candidates offer a promising starting point for developing broad-spectrum Cas13b-based antivirals against Dengue, with potential applicability to other RNA viruses.

Future efforts would focus on experimental validation of top candidates, expansion of the training dataset to improve model performance, and extension of the pipeline to other RNA viruses such as Chikungunya, and SARS-CoV-2 relevant to public health.

## Acknowledgment

We extend our sincere gratitude to Dr. Syed Faraz Ahmed, Department of Electrical and Electronic Engineering, University of Melbourne, for his careful review of this manuscript, constructive suggestions for improvement, and invaluable support in data curation and analysis.

## VII. Supplementary Material

**TABLE IV.**
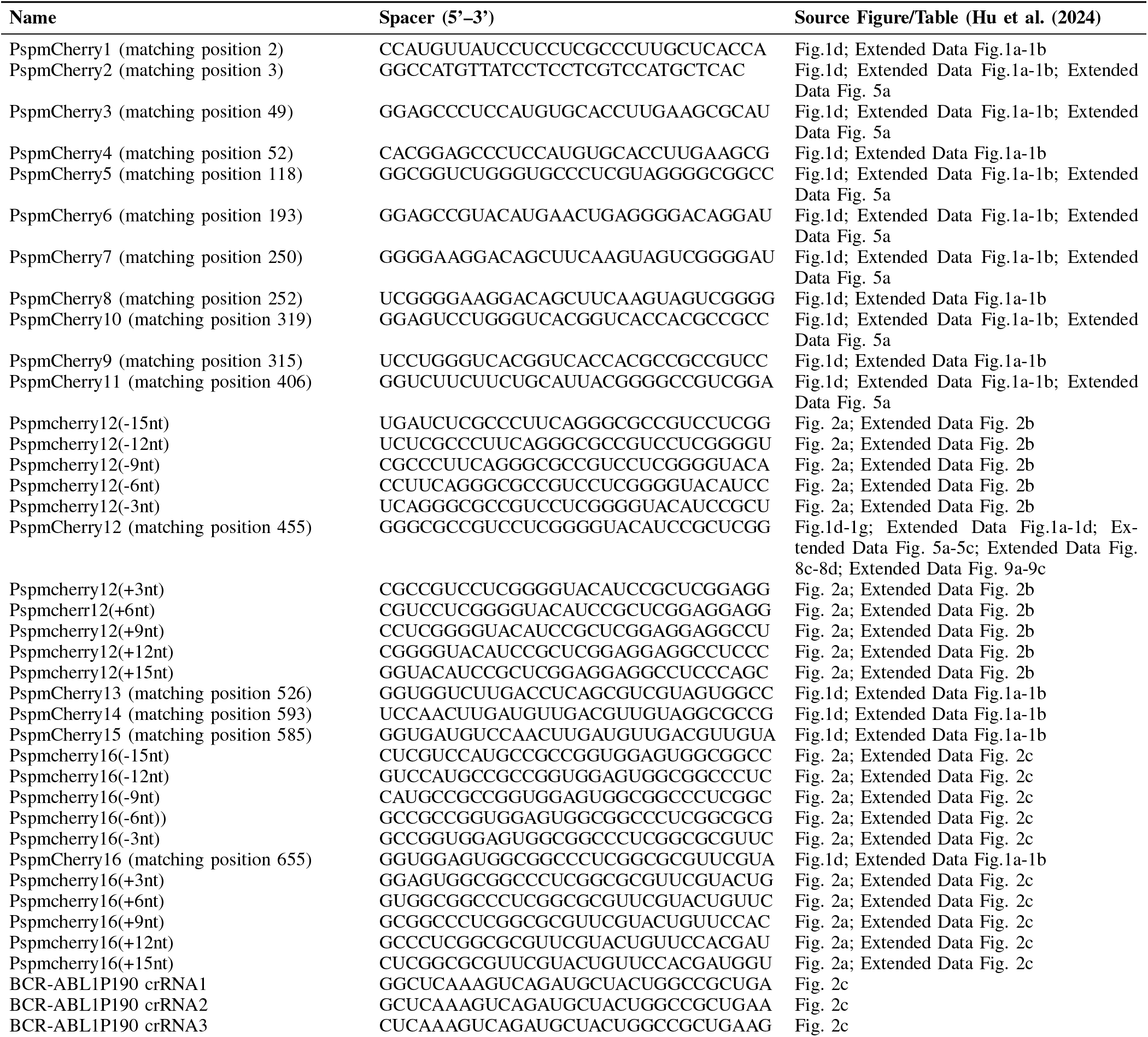
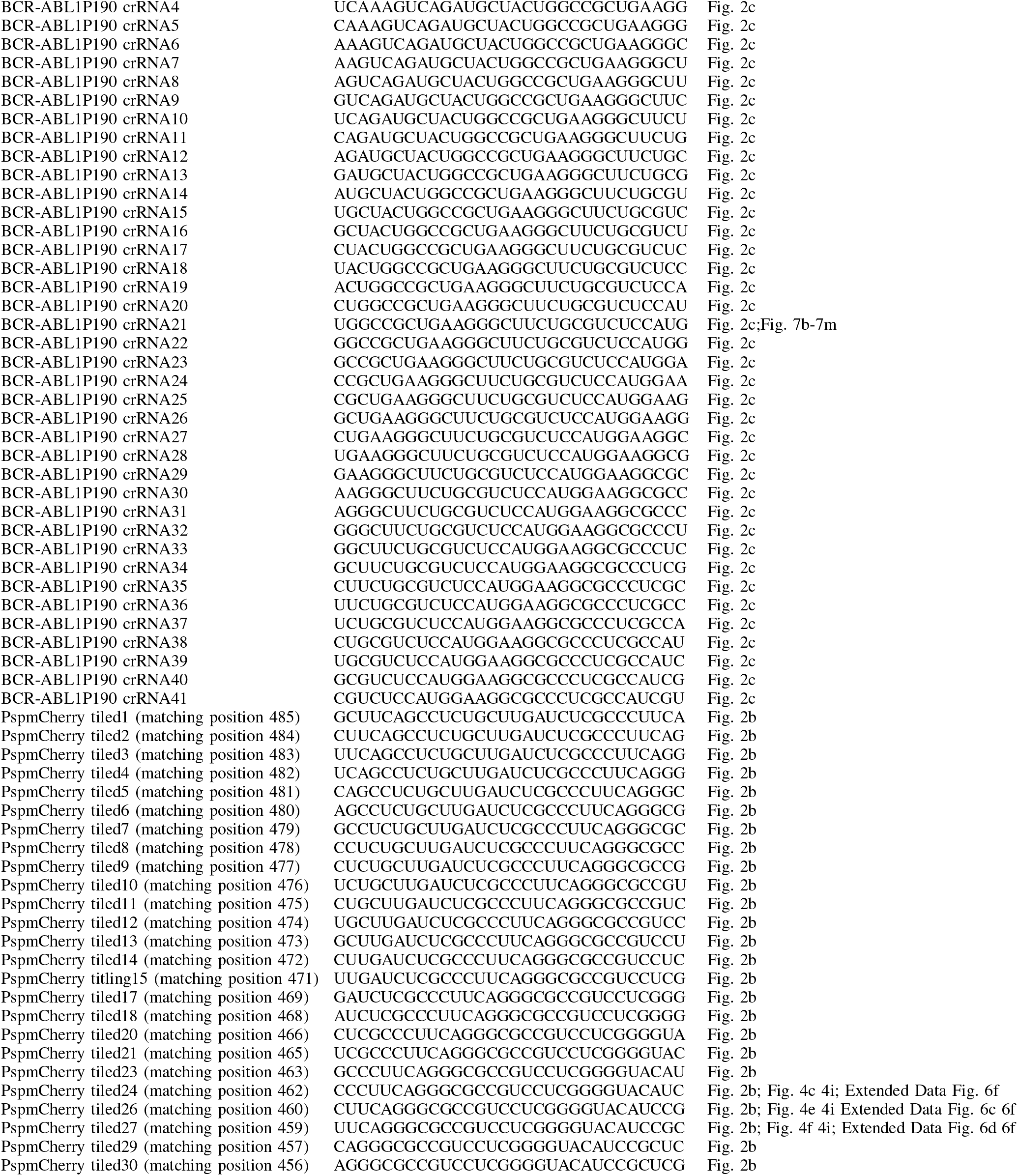
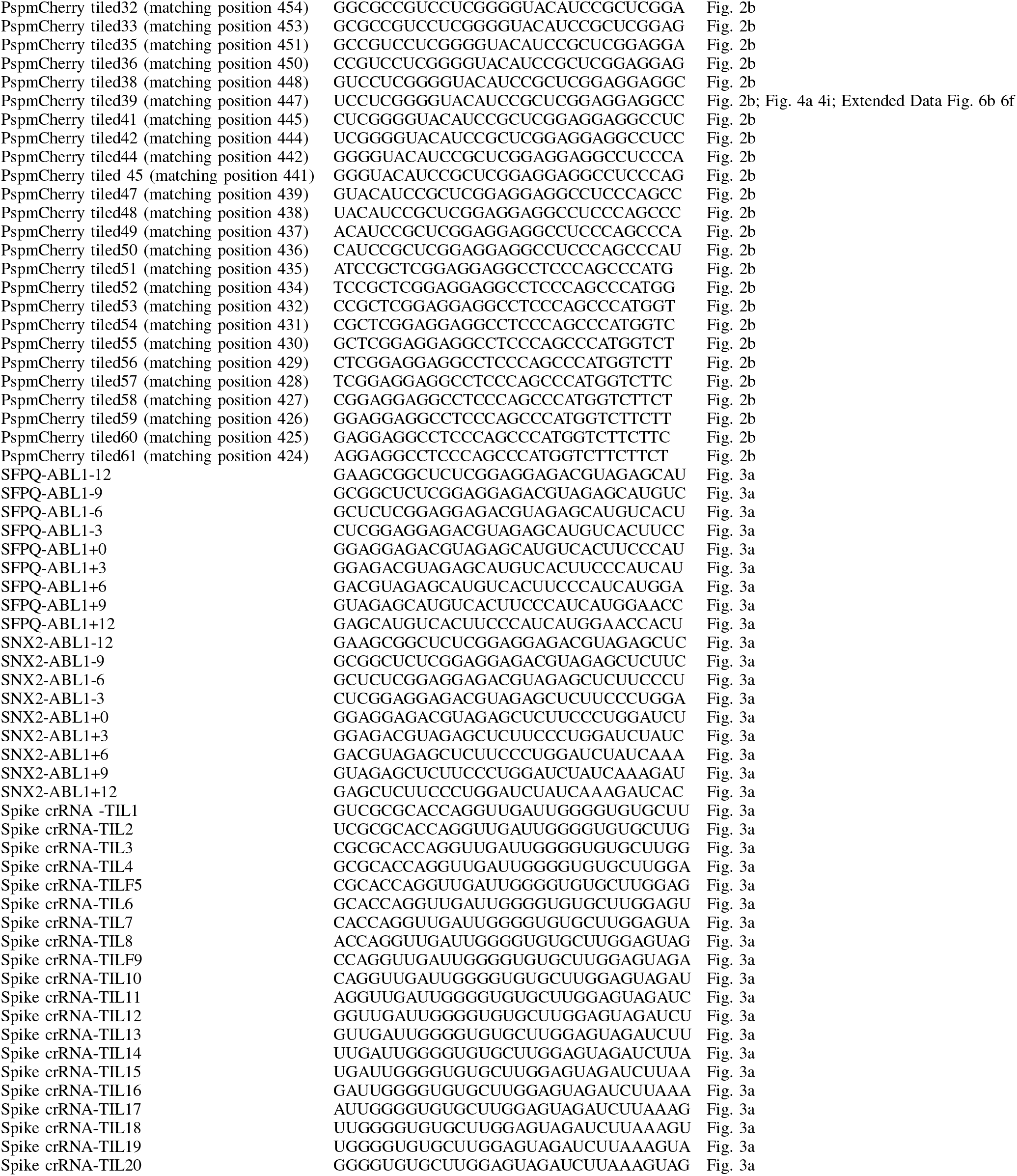
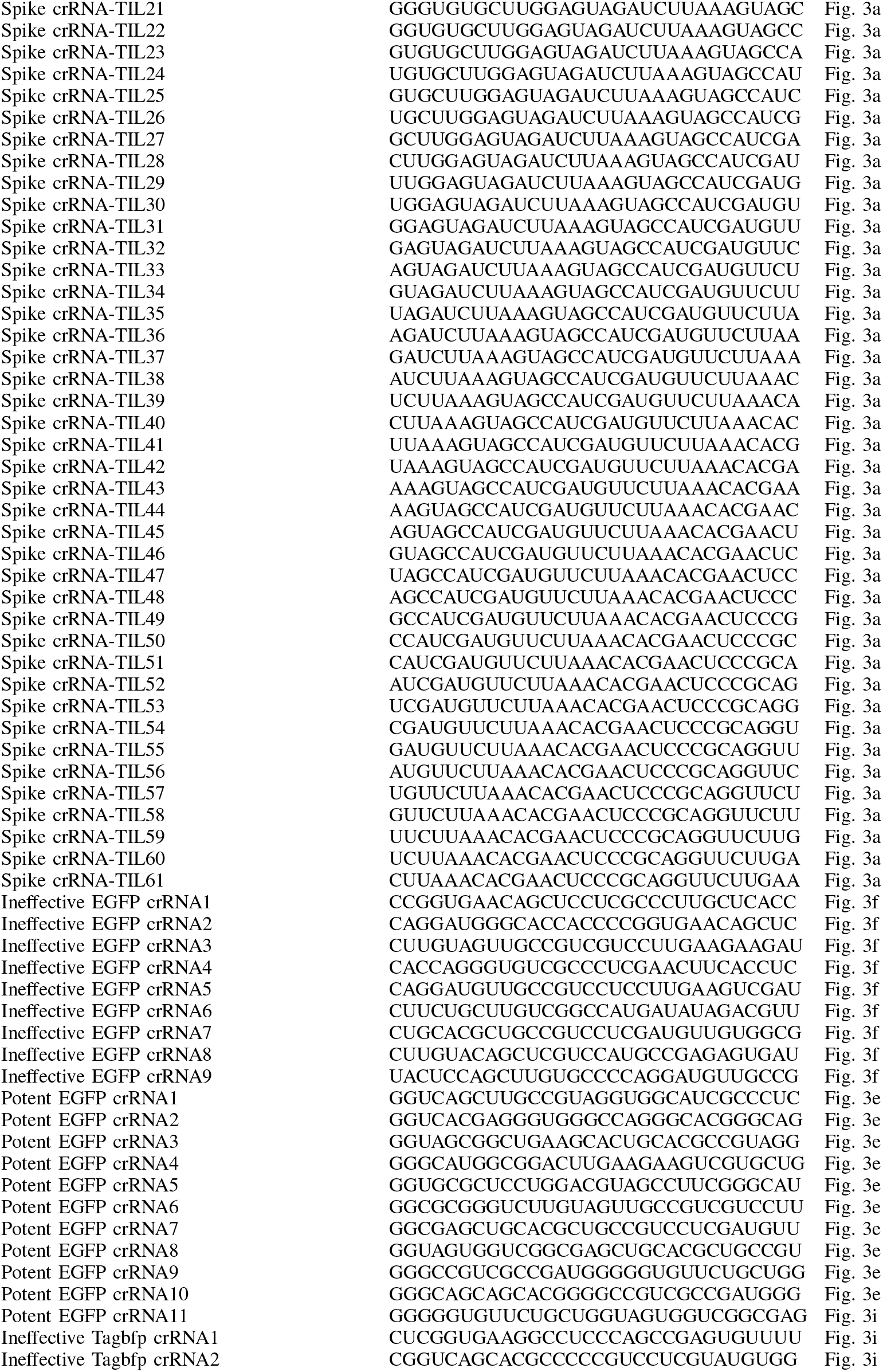
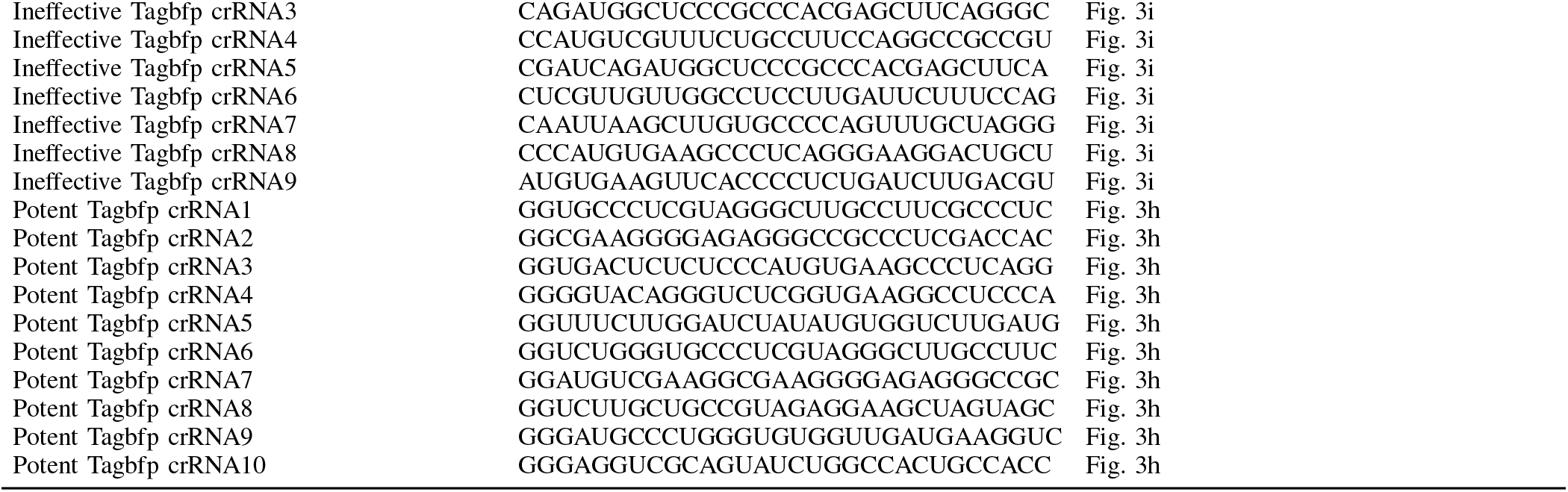
List of 245 crRNA spacers used to train the Cas13b efficiency models obtained from source data of [11]. The spacers can be found in the supplementary table 2 of [11] which lists the spacers along with the figures reporting their experimental efficacy values.

**TABLE V.** Summary of 157 Engineered Features used in ML Model.

| Category | Feature Description | Count |
| --- | --- | --- |
| **Positional One-Hot** | Binary indicator for A, C, G, T at each of 30 positions | 120 |
| **Nucleotide Counts** | Total count of A, C, G, T in spacer | 4 |
| **GC Content** | Percentage of G+C nucleotides | 1 |
| **Dinucleotides** | Frequency of all 16 pairs (AA, AC, ..., TT) | 16 |
| **Trinucleotides** | Frequency of key trimers (e.g., GGG, CCC) | 10 |
| **Motif: 5' End** | 'GG' at positions 1-2 (Binary) | 1 |
| **Motif: Penalties** | Count of 'C' at positions 1-4 | 1 |
|  | Count of 'C' at positions 11-17 (central region) | 1 |
| **Thermodynamics** | Minimum Free Energy (MFE) | 1 |
| **Homopolymers** | Presence of >3 consecutive identical bases | 1 |
| **Conservation** | Pre-computed conservation score | 1 |
| **Total** |  | **157** |

**TABLE VI.**
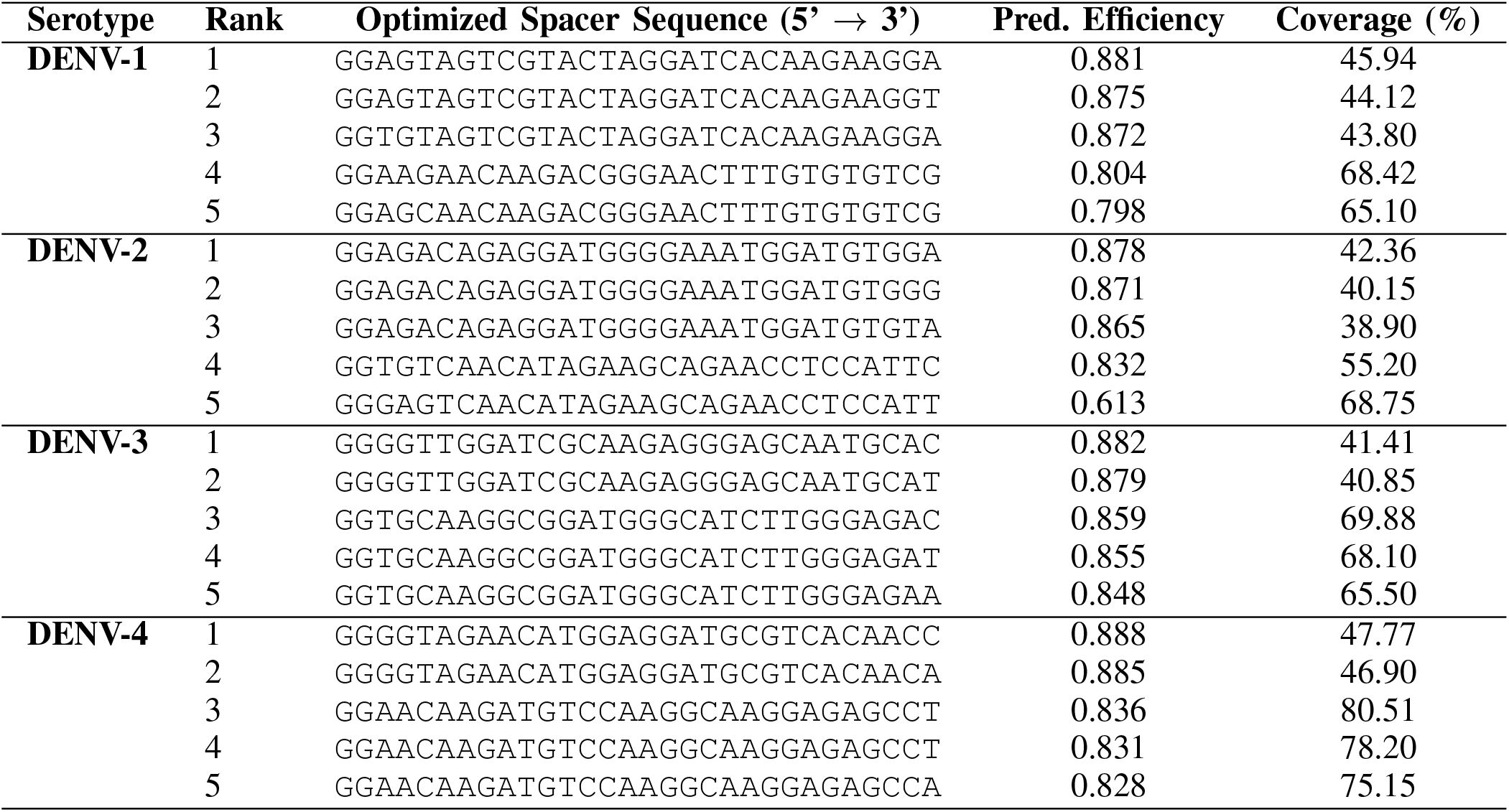
Top 5 Candidate crRNAs for DENV-1, DENV-2, DENV-3, and DENV-4.

